# Lianas in tropical dry forests have higher embolism resistance but similar hydraulic efficiency than lianas in rainforests

**DOI:** 10.1101/2023.02.01.526653

**Authors:** Caian S. Gerolamo, Luciano Pereira, Flavia R. C. Costa, Steven Jansen, Veronica Angyalossy, Anselmo Nogueira

## Abstract

Lianas are increasing in relative abundance and biomass, mainly in seasonally dry forests, but it is unclear if this is associated with their hydraulic strategy. Here, we ask whether liana of seasonally dry forests are safer and more efficient in water transport than those of rainforest, which could explain liana distribution patterns and their recent increases. We measured hydraulic traits on five pairs of congeneric liana species (tribe Bignonieae) on one seasonal dry Atlantic forest and one Amazon rainforest. The predawn and minimum water potential, and the water potential at which 50% of the maximum gas amount was discharged were, on average, more negative in the liana species of the seasonal forest. However, these patterns were not constant at the genus level. The positive hydraulic safety margins and hydraulic efficiency were similar among species congeners across sites. The Bignonieae lianas studied likely experience equally low levels of embolism during drought, and maintain a high conductive capacity with efficient use of xylem space, which may favor survival and growth across tropical forests. The likely evolutionary convergence of high hydraulic safety associated with the opportunistic strategy of rapid growth, especially in disturbed areas can favor the abundant liana species in seasonal forests.

**Highlight:** Tropical forest liana species have high hydraulic efficiency and high interspecific variability in hydraulic safety. Despite this variability, some seasonal forest liana species have greater hydraulic safety than rainforest lianas, indicating an evolutionary convergence across lineages.

## Introduction

Over the last decades, climatic seasonality intensified due to increased temperature and reduced precipitation in many tropical forests, increasing extreme events of drought (Williams *et al.*, 2007; Phillips *et al.*, 2010; Sherwood and Fu, 2014; Esteban *et al.*, 2021). In many tropical forests, there has been an increase in drought-induced tree mortality associated with the failure of the hydraulic transport system (Allen *et al.*, 2010; Anderegg *et al.*, 2016; Choat *et al.*, 2018). At the same time, lianas have been increasing in abundance in some tropical forests as seasonality increases (DeWalt *et al.*, 2015; Schnitzer, 2018; Schnitzer and van der Heijden, 2019; Medina-Vega *et al.*, 2021). For a plant to succeed in these new climatic conditions, it must resist drought and continue growing despite the drought effects. It has been hypothesized that the liana increase in seasonal dry tropical forests may derive from advantages associated with their hydraulic traits (Allen *et al.*, 2010; Anderegg *et al.*, 2016; Choat *et al.*, 2018). Therefore, it is fundamental to investigate the hydraulic features of lianas in forests with contrasting hydrological regimes and understand how this compares to the variation of lianas and other angiosperm growth forms across the globe.

Resilience of plant species to drought varies among environments and species, and can be related to xylem embolism resistance (Anderegg *et al.*, 2016). Drought induces more negative xylem water potentials, which may lead to the formation of air bubbles inside conductive cells and further propagation through “air-seeding” to neighboring conduits, decreasing the capacity to transport water, and potentially leading to complete hydraulic failure (Tyree and Ewers, 1991; Tyree and Zimmermann, 2002). Xylem embolism resistance is commonly evaluated through the water potential values at which the plant loses 50% (P50) or 88% (P88) of its conductive capacity by embolism (Tyree and Ewers, 1991; Anderegg *et al.*, 2016). This hydraulic trait describes the lethal threshold of dehydration in many plants and their susceptibility to hydraulic failure, with implications for productivity and survival in different ecosystems (Brodribb, 2009; Choat *et al.*, 2012; Urli *et al.*, 2013). P50 is positively associated with the plant’s minimum water potential throughout the day in the field, usually at midday and in the driest season (P_min_). Species that grow in drier environments attain more negative minimum water potentials in the xylem and are generally more resistant to drought-induced embolism than species that grow in moist environments (Choat *et al.*, 2012; Delzon and Cochard, 2014; Oliveira *et al.*, 2019). The P_min_ associated with P50 or P88 is a threshold for calculating the hydraulic safety margin (HSM = P_min_ – P50), indicating how much embolism the plant experiences in the field (Tyree and Ewers, 1991; Tyree *et al.*, 1994;

Meinzer *et al.*, 2009). Based on a global meta-analysis, there is evidence of a worldwide convergence in the hydraulic safety margin of angiosperm and gymnosperm trees, indicating a similar narrow HSM value (<1MPa) for tropical seasonally dry forests and tropical rainforests (Choat *et al.*, 2012). However, these hydraulic safety values come from different methods, and currently, we know that some earlier P50 data may be overestimated by measuring artefacts (Cochard *et al.*, 2013; Wheeler *et al.*, 2013; Torres-Ruiz *et al.*, 2014), reducing the hydraulic safety margin. Recent evidence showed that angiosperm trees in central and eastern Amazon forests operated far from their hydraulic limits when exposed to extreme natural drought, thereby exhibiting a hydraulic conservative strategy (Brum *et al.*, 2019; Garcia *et al.*, 2021).

In previous studies on embolism resistance, lianas from seasonally dry and wet tropical forests in Panama and China have similar or higher hydraulic vulnerability compared to co-occurring tree species (Zhu and Cao, 2009; De Guzman *et al.*, 2017; Zhu *et al.*, 2017; Van der Sande *et al.*, 2019). If so, lianas in drier environments would likely have compensating strategies to deal with water scarcity, such as greater stomatal control or increased leaf-shedding to reduce water loss, and/or access to deep water sources through deep roots (Schnitzer, 2018; Mantova *et al.*, 2021). On the other hand, liana species that grow in drier climates, such as semi-arid areas in Brazil (Carvalho *et al.*, 2016), have higher hydraulic safety than lianas of wet tropical forests in China and Panama (Zhu and Cao, 2009; Van der Sande *et al.*, 2013, 2019; Medina-Vega *et al.*, 2021), although there is also a notable interspecific difference. Thus, we hypothesize that seasonally dry tropical forests with lower availability of water may have lianas species that are more embolism resistant than liana species from wet rainforests, as has been shown for many tree species (Trueba *et al.*, 2017; Guillemot *et al.*, 2022). Consequently, if liana species from wet rainforests show reduced hydraulic safety and safety margin, they should be more vulnerable and prone to drought-induced mortality than lianas from seasonally dry forests.

Hydraulic safety is weakly related to the water transport efficiency in trees and shrubs (Gleason *et al.*, 2016, but see Wheeler *et al.*, 2005), and lianas tend to follow this weak or non-existing safety-efficiency trade-off (Zhu and Cao, 2009; Zhu *et al.*, 2017; Van der Sande *et al.*, 2013, 2019). Xylem hydraulic efficiency, measured as potential xylem-specific hydraulic conductivity, has been reported as a xylem space-use efficiency property (Bittencourt *et al.*, 2016). For a given stem diameter and length, a high hydraulic efficiency can supply a large leaf area at a lower energy cost, as long as compartmentalization is ignored (Bittencourt *et al.*, 2016). Regardless of the environment, there is extensive evidence that lianas have higher conductive efficiency and lower mechanical demand for support compared to tree species (Zhu and Cao, 2009; Zhu *et al.*, 2017; Van der Sande *et al.*, 2019; Medina-Vega *et al.*, 2021). The anatomical features of conducting cells of most lianas include a combination of wide and long vessels with narrow and short vessels, and tracheids (Carlquist, 1991; Tyree and Ewers, 1991; Angyalossy *et al.*, 2012, 2015; Gerolamo and Angyalossy, 2017), favoring a higher xylem space-use efficiency and hydraulic conductivity compared to trees (Ellmore and Ewers, 1985; Ewers and Fisher, 1989; Tyree and Zimmermann, 2002). High hydraulic efficiency can be considered as an essential feature of the liana’s success since greater conductivity increases the delivery of water to plant tissues, and may favor photosynthesis and consequently the stem and leaf area growth (Zhu and Cao, 2009; Van der Sande *et al.*, 2013, 2019). However, if liana species of seasonal forest lianas have increased in abundance then perhaps they can have a higher xylem hydraulic efficiency compared to liana species from wet forests.

Hydraulic safety and efficiency integrate critical aspects of plant physiology, structure, and their interaction with the microclimate and soil (Choat *et al.*, 2012; Delzon and Cochard, 2014; Skelton *et al.*, 2015; Fontes *et al.*, 2020). These hydraulic features are associated with niche differentiation across environmental gradients (Engelbrecht *et al.*,2007; Brodribb, 2017; Brum *et al.*, 2019; Oliveira *et al.*, 2019), and are rarely explored on liana assemblages across contrasting forest types (Medina-Vega *et al.*, 2021). Therefore, in this study, we investigated the hydraulic safety and efficiency of five pairs of congeneric liana species of the tribe Bignonieae distributed in two distinct Neotropical forest types: the wet Amazon rainforest (~ 2 dry months) and seasonal semideciduous Atlantic forest (~6 dry months). The tribe Bignonieae (Bignoniaceae) is the most diverse lineage of lianas in the Neotropics (Gentry, 1991; Lohmann *et al.*, 2013). The most parsimonious interpretation of the biogeographic history of this group indicates that the crown node of Bignonieae originated in South American rainforests, diversifying in the lowland Amazon forest, and more recently spread to seasonally drier forests and savannas (Lohmann *et al.*, 2013). For lianas to have survived in seasonally drier forests, they must resist drought. After and along this period, the high conductive efficiency would favor the growth of lianas. In this context, the main question that emerges is whether liana species of dry seasonal forests are safer and more efficient in transporting water than liana species of wet rainforests. We hypothesized that: (H1) liana species of dry seasonal forest cope with a lower availability of water and consequently have a higher hydraulic safety compared to liana species of wet rainforest, minimizing the risk of hydraulic failure under drought; (H2) liana species of dry seasonal forest have a more efficient xylem hydraulic transport than liana species of wet rainforest, not supporting the safety-efficiency trade-off. If the safety-efficiency pattern between forest types occurs in different genera of lianas, the recurrent hydraulic features of liana species in the dry seasonal forest associate with the biogeographic history of the clade Bignonieae would indicate an evolutionary convergence under drier conditions. Furthermore, by combining a large plant hydraulic safety dataset, we can evaluate the variation in hydraulic safety of lianas concerning different forest types and plant habits within a global context, and test (H3) if liana species of dry seasonal forests have higher hydraulic safety than trees, shrubs, and lianas from other tropical forests. Addressing these hypotheses and understanding how hydraulic safety and efficiency changes in lianas from different tropical forests allow us to provide a valuable framework for mechanistic models of plant hydraulics under climate change.

## Materials and Methods

### Study site and climate

The study was carried out in two contrasting forest sites marked by the differentiation of the water regimes. The first one was a tropical wet rainforest site located at Ducke Reserve, a 10,000 ha reserve (WF) situated in the central region of the Amazon basin, Manaus, Brazil (02°55’S, 59°58’W). The second was a drier seasonal semideciduous forest site located at Santa Genebra Reserve, a 250 ha reserve (SF) in the Atlantic Forest domain in São Paulo, Brazil (22°45’ S, 47°33’W).

According to the Koppen-Geiger climate classification, the Ducke Reserve experiences a tropical ‘Am’ climate, with dry and rainy seasons ruled by monsoons (Peel *et al.*, 2007). Annual rainfall and average maximum temperature in the 2018 collection were 2147 mm and 32 °C, respectively, with August being one of the driest months with 86 mm of precipitation (Supplementary Fig. S1 - Climate diagram; data obtained from the Station of Ducke Reserve, Laboratory of Climatic Modeling of National Institute for Amazonian Research managed by Dr. Luiz A. Candido). The vegetation of the Ducke Reserve was an old-growth perennial rainforest with a high diversity of tree and liana species. The forest has a closed canopy of 30-37 m and emerging trees reaching 45 m (Guillaumet and Kahn, 1982; Ribeiro *et al.*, 1999; Rocha *et al.*, 2022). The topography was heterogeneous, with elevations ranging from 40 to 140 m above sea level (Ribeiro *et al.*, 1999). Soils form a continuum of clayey oxisols on plateaus, with increasing sand on the slopes, until they become pure sand at the bottom of the valleys, forming small floodplains (Chauvel *et al.*, 1987; Mertens, 2004).

According to the Koeppen-Geiger climate classification Campo, Santa Genebra Reserve experiences a tropical ‘Cwa’ climate (Peel *et al.*, 2007). Annual rainfall and average maximum temperature in 2020 were 871 mm and 40 °C, respectively, with September being a dry month at the end of the dry season with 18 mm of precipitation (Supplementary Fig. S1; data obtained from the http://clima.iac.sp.gov.br/ in Campinas station). The vegetation of the Santa Genebra Reserve was a seasonal semideciduous forest, occupying 85% of the reserve, with a high diversity of tree and lianas species and an almost continuous canopy at about 15 m in height with emerging trees reaching 30 m (Morellato, 1991; Morellato and Leitao-Filho, 1996). The topography was homogeneous, with elevations ranging from 580 to 610 m above sea level, and soils were typical dystrophic red argisol (Mendes *et al.*, 2013).

### Species collection and design

We selected five abundant species of lianas in each forest type, totaling ten species (Supplementary Table S1). These species were chosen for being congeneric species pairs across forest types. One species was from the Amazon rainforest site, and the other was from the drier seasonal Atlantic forest site. In addition, these species were abundant in these forests (Morellato, 1991; Morellato and Leitao-Filho, 1996; Gerolamo *et al.*, 2022). We used congeneric species pairs to assess whether habitat-associated drought traits repeatedly occurred in different phylogenetic lineages. In our sampling, we included only mature individuals, i.e., plants that reached the forest canopy and had a stem diameter >1 cm at 1.30 m from the rooting point. Six to eight individuals were marked and sampled for each species. Each liana was located more than 10 meters from each other without any stem connection with other lianas to avoid clonal plants (Schnitzer, 2006). The distribution of marked individuals/species in each forest varied according to the habitat. At Ducke reserve, we found two species of lianas predominantly distributed in the valleys close to the watercourse. The other three species were located predominantly on the plateaus and slopes (Gerolamo *et al.*,2022). At Santa Genebra reserve, the five species occurred in the plateaus and close to the main trails in clearing areas. The liana species of wet rainforest and seasonal forest were collected in the dry season in 2018 and 2020, respectively (Supplementary Fig. S1). After field collection, the branches were taken to the laboratory to estimate the vulnerability to embolism, as described below. A voucher for each species was deposited at the Herbarium of the University of São Paulo (SPF; the acronym of (Thiers, 2017) (Supplementary Table S1).

### Vulnerability to xylem embolism

We collected a branch (ca. 2 m in length) of six to eight individuals longer than the maximum vessel length for each species. The maximum vessel lengths were measured using an air-injection method in a minimum of three individuals per species (Greenidge, 1952; Ewers and Fisher, 1989; Jacobsen *et al.*, 2012), and values were in the range of 56-92 cm. We collected about three branches per day in the early morning between 5:30 am, and 7:30 am local time (Manaus and Campinas, Brazil) to avoid artificial embolism propagation by cutting stems with rather negative xylem water potentials. All branches were cut from the canopy, and each branch was sprayed with water, placed inside dark plastic bags with moistened paper towels, and the cut ends were immersed in a beaker with water to prevent desiccation. We transported the branches to the laboratory within ca. 35 min. They were kept under dark plastic bags with moistened paper towels for at least one hour before starting the vulnerability curves. The hydraulic vulnerability curves were constructed using the manual pneumatic method (Pereira *et al.*, 2016; Bittencourt *et al.*, 2018; Zhang *et al.*, 2018). The total air discharging tube volume of 3.915 ml was calculated from the tubing datasheets, in which each rigid tubing (Cole-Parmer, USA) had 1.304 ml (for details, see Bittencourt *et al.*, 2018). The air volume was estimated from the amount of gas extracted over 2.5 min from the cut end of the branch after applying a partial vacuum (~ 50 kPa absolute pressure). The measurements were carried out before the development of an automated pneumatic apparatus (Pereira *et al.*, 2020) and estimation of the best extraction duration (Paligi *et al.*, 2021; Yang *et al.*, 2021), but we expect a relatively low error for the P50 estimation considering the good agreement between the manual pneumatic method and other methods (Brum *et al.*, 2023).

Branches were bench dehydrated, and the xylem water potential was measured using a pressure chamber (PMS 1000; PMS Instruments Co., Albany, OR, USA; (Scholander *et al.*,1965). The branches were bagged for 30 min to 1 hour to balance the leaf and xylem water potential, and then two leaves from each branch were used to estimate the average xylem water potential in each measurement. About 8 to 10 measurements were conducted on each branch. The last measurement was taken when the percentage of air discharge had stabilized, the leaves were completely dehydrated, and/or the xylem water potential exceeded −8 to −9 MPa. We pooled the data for each branch and fitted a sigmoidal curve to the data, relating the percentage of gas discharged (PAD) to the xylem water potential. In the sigmoidal curve, P50 and slope *(b)* were the fitted parameters (Pammenter and Willigen, 1998) and P88 was predicted from the fitted model: PAD = 100/ {1 + exp [*b* (Ψ– P50)]}. We fitted the curves at the individual level and then pooled individuals’ curves by species. P50 and P88 were the xylem water potential at which 50% and 88% of the maximum gas amount were discharged, respectively. We used both P50 and P88 as indicators of xylem resistance to hydraulic failure.

### Predawn and midday water potential

Leaf water potential (MPa) was measured at predawn and midday and during the peak of the dry season in each forest (August-September; Supplementary Fig. S1) using a pressure chamber. About three to six individuals of each species were selected (same individuals as used for the vulnerability curves), and leaves of two to three individuals per day were measured. For each chosen liana, three fully developed mature canopy leaves per individual were collected during the morning between 5:30 am, and 7:00 am (P_predawn_). Another three leaves were collected between 11:30 am and 2:00 pm (P_midday_). The leaf water potential was measured immediately after collecting the leaves in the field. At the whole-plant level, the leaves were more exposed to the low water potentials of the atmosphere and consequently experienced more negative water potentials than those found in the stem (Tyree, 1988; Guan *et al.*, 2021). Due to the stem-leaf transition of the water potential gradient, if the stomata were not closed, we selected the most positive value of the three leaves measured in P_midday_ as a reference to represent the minimum water potential of the stem xylem (P_min_). The xylem hydraulic safety margin (HSM) was calculated as HSM = P_min_ - P50, where P_min_ was the minimum seasonal leaf water potential measured in the field during the dry season (August-September, Supplementary Fig. S1).

### Anatomical measurements

In order to establish a standard for the distance from the apex, which was necessary for comparison of the vessel diameter (Gasson and Baas, 1983; Rosell and Olson, 2014), we sectioned samples from the base of the stem (about 1 meter from the apex) from the same branches that were used for the vulnerability curves to perform the anatomical procedures. Each stem sample was fixed in FAA (formaldehyde, acetic acid, and 50% ethanol, Johansen, 1940) and stored in 70% ethanol. The samples, composed of bark (secondary phloem and periderm) and secondary xylem, were embedded in polyethylene glycol 1500 (PEG-1500, (Rupp, 1964).

Transverse, longitudinal, radial, and tangential sections of each sample were made with a sliding microtome, following Barbosa et al. (2010). Sections were stained in 1% astra blue, and 1% safranin (Bukatsch, 1972; Kraus and Arduin, 1997), and permanent slides were mounted for anatomical analyses. To estimate vessel diameter and vessel frequency, as well as the theoretical specific hydraulic conductivity, we photographed each histological slide with a photomicroscope (Leica DML and camera DFC 310FX). The images were analyzed using ImageJ version 1.45d software (National Institutes of Health, Bethesda, MD, USA; http://rsb.info.nih.gov/ij/). We used the procedures established by (Scholz *et al.*, 2013), measuring at least 100 vessels per stem sample to characterize vessel diameter and counting the number of vessels in four areas of 1 mm^2^ to estimate the vessel frequency per sample. All anatomical measurements were accomplished in the interwedge regions, i.e., portions of the stem that have regular secondary growth in Bignonieae lianas (Gerolamo and Angyalossy, 2017).

### Specific hydraulic conductivity

Theoretical specific hydraulic conductivity (Ks; Kg.m.Mpa^−1^.s^−1^) was used to indirectly describe water transport efficiency. Using vessel diameter and vessel frequency of each individual, we calculated the Ks (Equation 1) following Hagen–Poiseuille’s law (Tyree and Ewers, 1991; Poorter *et al.*, 2010), as

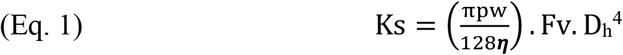

where pw was the density of water at 20°C (998.2 kg.m^−3^), ***η*** was the water viscosity at 20 °C (1.002 10^−3^ Pa.s^−1^), Fv was the vessel frequency per mm^2^, and Dh was the hydraulic vessel diameter (m).

We calculated D_h_ (Equation 2) considering both wide and narrow vessels, as

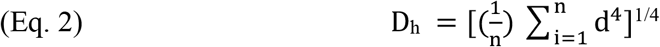

where n was the number of vessels and d was the vessel diameter.

Narrow vessels could be misinterpreted as tracheids (Carlquist, 1985). However, we analyzed the dissociated xylem cells in macerated tissue and did not find any tracheid in all liana species sampled. Ks was assumed to be higher than actual conductivity values because the resistance of the end walls of vessels (Gibson *et al.*, 1985; Sperry *et al.*, 2008), and embolized vessels were not taken into account. Here, we assumed that these additional resistances did not change the relative differences found in Ks between species, since Ks would scale positively with actual conductivity values (Ewers *et al.*, 1989; Tyree and Zimmermann, 2002).

### Data analysis

Following the first hypothesis (H1), we expected that liana species from the dry seasonal forest cope with a lower availability of water (lower P_predawn_ and P_min_) and consequently had a higher hydraulic safety (lower P50) compared to liana species of wet rainforest, minimizing the risk of hydraulic failure (HSM) under drought. To evaluate our H1, we assessed the effect of forest type and genus (predictor variables) on predawn water potential (P_predawn_), minimum seasonal water potential (P_min_), hydraulic safety (P50), and safety margin (HSM) (response variables), using a model selection approach for each response variable (Legendre and Legendre, 1998), described below. We built linear mixed models (LMM) with Gaussian error distribution for each response variable, considering the forest type and genus as fixed factors and the species nested within a genus as a random factor on the intercept. A random factor was defined by the sampling design to account for the nonindependence of individuals nested in species and species nested by genus.

Following the second hypothesis (H2), we expected that liana species of dry seasonal forest had a more efficient xylem hydraulic transport (Ks) than liana species of wet rainforest, not supporting the safety-efficiency trade-off. To evaluate our H2, we used the same model selection approach, evaluating the effect of forest type and genus on the theoretical specific hydraulic conductivity (Ks). Also, we tested the relationship between theoretical specific hydraulic conductivity and hydraulic safety (P50) to investigate the safety-efficiency trade-off across plants directly. We included the random term in the models above to describe the nested structure of our sampling design. Data of Ks were log-transformed before analysis to achieve normality.

The models used to test the H1 and H2 included the genera as a fixed factor representing the different lineages of lianas within the tribe Bignonieae. In this case, a concordant pattern of hydraulic features could emerge of differences between forest types across genera. If so, we can infer a pattern of convergence of hydraulic features in the liana species occurring in the dry seasonal forest due to the particular biogeographic history of this plant group – the most recent occupation of lianas in drier forests and savannas independently in each genus investigated.

In all cases above, we evaluated the most suitable minimal model from a set of models that included a null model (only intercept without forest type or genus as fixed factors). The minimum suitable model had the lowest Akaike information criterion (AIC) value, and models with ΔAIC< 2 were considered plausibly similar. We validated the selected models by visually verifying the homogeneity of variance and normality of the residuals. For all statistical analyses, we used R v.3.3.0 (R Core Team, 2020) with base packages and the lmer function of lme4 (Bates, 2010).

To contextualize the variation in hydraulic safety of lianas in a global framework and to test the third hypothesis (H3) that liana species of dry seasonal forests were hydraulically safer than shrubs, trees, and lianas from other tropical forests, we combined a large dataset with average hydraulic safety (P50) for angiosperm species, including trees, shrubs, succulents, herbs and lianas from different forest sites (Zhu and Cao, 2009; Choat *et al.*,2012; Carvalho *et al.*, 2016; Chen *et al.*, 2017, 2021; Oliveira *et al.*, 2019; Tan *et al.*, 2020; De Guzman *et al.*, 2021; Medina-Vega *et al.*, 2021; Smith-Martin *et al.*, 2022). Data on the number of species per habit included in each study can be viewed in the Supplementary data Table S2. We did not include studies that used air injection and centrifugation techniques to estimate P50 because we know that these techniques can overestimate P50 values due to effervescence (Yin and Cai, 2018) and open-vessel artefacts (Cochard *et al.*, 2013; Wheeler *et al.*, 2013; Torres-Ruiz *et al.*, 2014; Pereira *et al.*, 2021). Data of tree, shrub and liana species of tropical rainforest and seasonal tropical forest were separated for analysis. We compared the hydraulic safety range (P50 max – P50 min) of lianas in the seasonal forest with the rainforest and trees and shrubs of other tropical forests. The difference in P50 values was tested using t-tests and α = 0.05.

## Results

The hydraulic traits of the five pairs of congeneric Bignonieae lianas differed unevenly between forest types (Table 1).

**Table 1.**
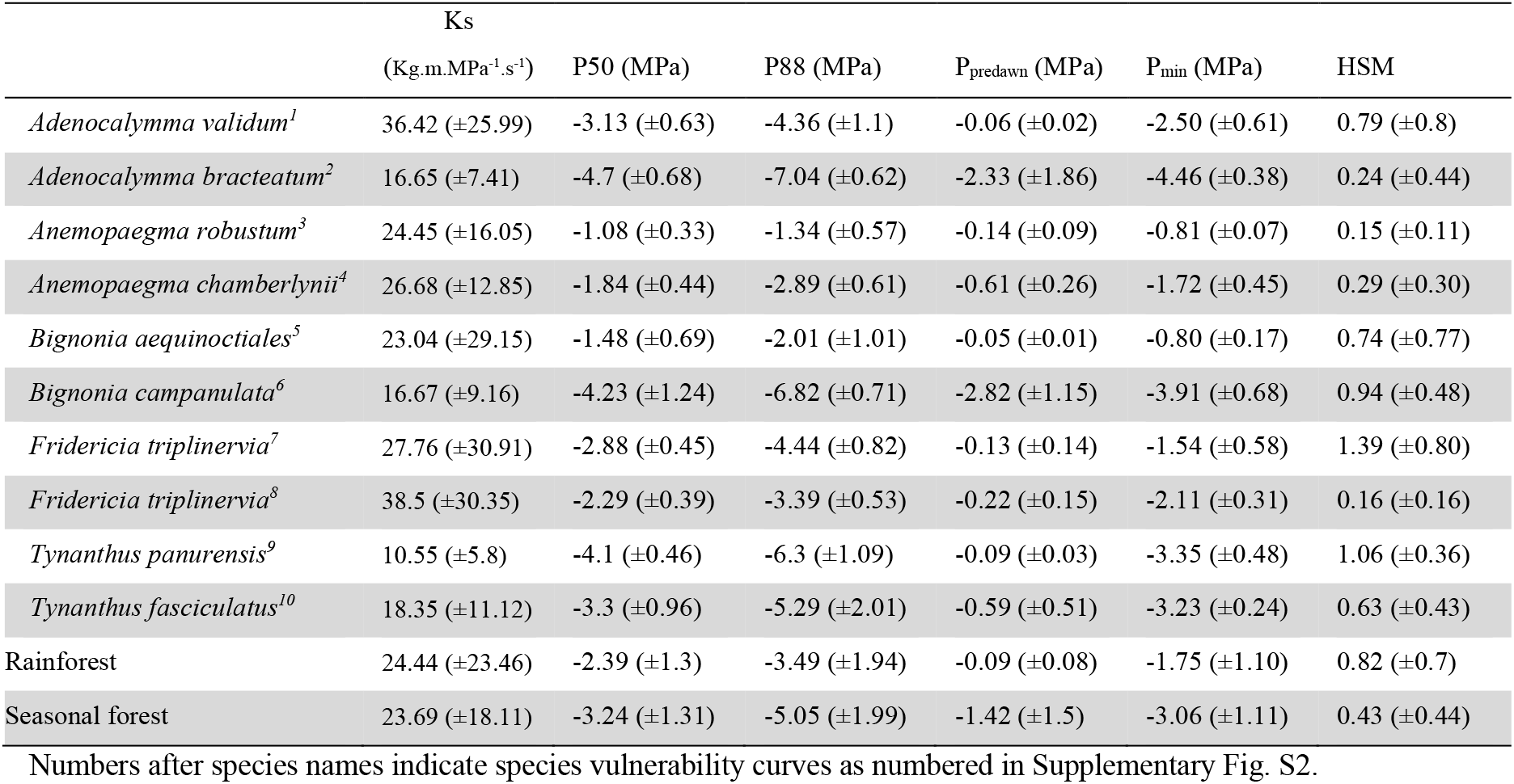
Mean value (±SD) of theoretical specific hydraulic conductivity (Ks), xylem pressure at which 50% and 88% of maximum gas amount was discharged (P50 and P88), predawn water potential (P_predawn_), minimum seasonal water potential (P_min_), hydraulic safety margin (HSM = P_min_ – P50) and, for congeneric liana species of Bignonieae occurring in two contrasting forests (rainforest and seasonal forest).

### Hydraulic safety and efficiency in lianas

Liana species of the seasonal forest had lower (more negative) predawn water potential (P_predawn_) values than congeneric liana species of the rainforest (Fig. 1A; Model 1 in Table 2). In the rainforest and seasonal forest, most species reached P_predwan_ close to zero, but *Adenocalymma* and *Bignonia* in the seasonal forest reached values close to −2.5 MPa (Fig. 1A; Table 1). The minimum seasonal water potential (P_min_) of liana species of the seasonal forest was 1.4 to 4.8 times more negative than congeneric rainforest lianas, except for *Tynanthus* (Fig. 1B; Table 1). However, the differences in the P_min_ averages between liana species of rainforest and seasonal forest were genus-dependent (Model 6 in Table 2). Three genera (*Adenocalymma*, *Anemopaegma*, and *Bignonia*) had more negative values of P_min_ in the seasonal forest. In contrast, the other two genera had more similar P_min_ values between the two forest types.

**Table 2.** Model selection results for each response variable: predawn water potential (P_predawn_), minimum seasonal water potential (P_min_), xylem pressure at which 50% of the maximum gas amount was discharged (P50), hydraulic safety margin (HSM), and theoretical specific hydraulic conductivity (Ks) of lianas for two forest types and five genera. Bold lines are the selected models for each response variable. Gaussian probability distribution was used in all mixed models, including none, one or two fixed factors - forest (categorical) and genus (categorical) - and a random term describing the nested sampling design. Degrees of freedom (df), Second-order Akaike’s information criterion (AICc) and ΔAICc (AICcmodel-AICcminimum) are also shown.

| Model | Response variable | Fixed factors | Random term | df | AICc | $\Delta\text{AICc}$ | Figure |
| --- | --- | --- | --- | --- | --- | --- | --- |
| **1** | **Log($P_{\text{predawn}}$)** | **Forest** | **1\|Genus/Species** | **5** | **121.5** | **0** | **1A** |
| 2 |  | Forest x Genus | 1\|Genus/Species | 13 | 124.0 | 2.4 |  |
| 3 |  | 1 | 1\|Genus/Species | 4 | 124.4 | 2.8 |  |
| 4 |  | Forest + Genus | 1\|Genus/Species | 9 | 124.5 | 3.0 |  |
| 5 |  | Genus | 1\|Genus/Species | 8 | 126.1 | 4.6 |  |
| **6** | **$P_{\min}$** | **Forest x Genus** | **1\|Genus/Species** | **13** | **99.2** | **0** | **1B** |
| 7 |  | Forest + Genus | 1\|Genus/Species | 9 | 100.8 | 1.6 |  |
| 8 |  | Forest | 1\|Genus/Species | 5 | 102.3 | 3.1 |  |
| 9 |  | Genus | 1\|Genus/Species | 8 | 102.6 | 3.3 |  |
| 10 |  | 1 | 1\|Genus/Species | 4 | 104.6 | 5.4 |  |
| **11** | **P50** | **Forest x Genus** | **1\|Genus/Species** | **13** | **191.4** | **0** | **1C** |
| 12 |  | Forest + Genus | 1\|Genus/Species | 9 | 196.8 | 5.3 |  |
| 13 |  | Genus | 1\|Genus/Species | 8 | 196.8 | 5.4 |  |
| 14 |  | Forest | 1\|Genus/Species | 5 | 199.4 | 8.0 |  |
| 15 |  | 1 | 1\|Genus/Species | 4 | 199.8 | 8.3 |  |
| **16** | **Safety Margin** | **1** | **1\|Genus/Species** | **4** | **102.5** | **0** | **1D** |
| 17 |  | Forest | 1\|Genus/Species | 5 | 103.7 | 1.2 |  |
| 18 |  | Genus | 1\|Genus/Species | 8 | 110.3 | 7.8 |  |
| 19 |  | Forest + Genus | 1\|Genus/Species | 9 | 112.1 | 9.6 |  |
| 20 |  | Forest x Genus | 1\|Genus/Species | 13 | 116.8 | 14.3 |  |
| **21** | **Log(Ks)** | **1** | **1\|Genus/Species** | **4** | **161.7** | **0.0** | **1F** |
| 22 |  | Forest | 1\|Genus/Species | 5 | 165.1 | 3.4 |  |
| 23 |  | Genus | 1\|Genus/Species | 8 | 168.3 | 6.7 |  |
| 24 |  | Forest + Genus | 1\|Genus/Species | 9 | 172.1 | 10.4 |  |
| 25 |  | Forest x Genus | 1\|Genus/Species | 32 | 177.4 | 15.7 |  |
| **26** | **Log(Ks)** | **1** | **1\|Genus/Species** | **4** | **159.4** | **0.3** | **2** |
| 27 |  | P50 | 1\|Genus/Species | 5 | 159.1 | 0 |  |
| 28 |  | P50 + Forest | 1\|Genus/Species | 6 | 160.4 | 1.3 |  |
| 29 |  | Forest | 1\|Genus/Species | 5 | 161.5 | 2.4 |  |
| 30 |  | P50 x Forest | 1\|Genus/Species | 7 | 163. | 3.8 |  |

**Table 2.** Model selection results for each response variable: predawn water potential ( $P_{\text{predawn}}$ ), minimum seasonal water potential ( $P_{\text{min}}$ ), xylem pressure at which 50% of the maximum gas amount was discharged ( $P_{50}$ ), hydraulic safety margin (HSM), and theoretical specific hydraulic conductivity ( $K_s$ ) of lianas for two forest types and five genera.
| Model | Response variable | Fixed factors | Random term | df | AICc | $\Delta\text{AICc}$ | Figure |
| --- | --- | --- | --- | --- | --- | --- | --- |
| **1** | **Log($P_{\text{predawn}}$)** | **Forest** | **1\|Genus/Species** | **5** | **121.5** | **0** | **1A** |
| 2 |  | Forest x Genus | 1\|Genus/Species | 13 | 124.0 | 2.4 |  |
| 3 |  | 1 | 1\|Genus/Species | 4 | 124.4 | 2.8 |  |
| 4 |  | Forest + Genus | 1\|Genus/Species | 9 | 124.5 | 3.0 |  |
| 5 |  | Genus | 1\|Genus/Species | 8 | 126.1 | 4.6 |  |
| **6** | **$P_{\text{min}}$** | **Forest x Genus** | **1\|Genus/Species** | **13** | **99.2** | **0** | **1B** |
| 7 |  | Forest + Genus | 1\|Genus/Species | 9 | 100.8 | 1.6 |  |
| 8 |  | Forest | 1\|Genus/Species | 5 | 102.3 | 3.1 |  |
| 9 |  | Genus | 1\|Genus/Species | 8 | 102.6 | 3.3 |  |
| 10 |  | 1 | 1\|Genus/Species | 4 | 104.6 | 5.4 |  |
| **11** | **$P_{50}$** | **Forest x Genus** | **1\|Genus/Species** | **13** | **191.4** | **0** | **1C** |
| 12 |  | Forest + Genus | 1\|Genus/Species | 9 | 196.8 | 5.3 |  |
| 13 |  | Genus | 1\|Genus/Species | 8 | 196.8 | 5.4 |  |
| 14 |  | Forest | 1\|Genus/Species | 5 | 199.4 | 8.0 |  |
| 15 |  | 1 | 1\|Genus/Species | 4 | 199.8 | 8.3 |  |
| **16** | **Safety Margin** | **1** | **1\|Genus/Species** | **4** | **102.5** | **0** | **1D** |
| 17 |  | Forest | 1\|Genus/Species | 5 | 103.7 | 1.2 |  |
| 18 |  | Genus | 1\|Genus/Species | 8 | 110.3 | 7.8 |  |
| 19 |  | Forest + Genus | 1\|Genus/Species | 9 | 112.1 | 9.6 |  |
| 20 |  | Forest x Genus | 1\|Genus/Species | 13 | 116.8 | 14.3 |  |
| **21** | **Log($K_s$)** | **1** | **1\|Genus/Species** | **4** | **161.7** | **0.0** | **1F** |
| 22 |  | Forest | 1\|Genus/Species | 5 | 165.1 | 3.4 |  |
| 23 |  | Genus | 1\|Genus/Species | 8 | 168.3 | 6.7 |  |
| 24 |  | Forest + Genus | 1\|Genus/Species | 9 | 172.1 | 10.4 |  |
| 25 |  | Forest x Genus | 1\|Genus/Species | 32 | 177.4 | 15.7 |  |
| **26** | **Log($K_s$)** | **1** | **1\|Genus/Species** | **4** | **159.4** | **0.3** | **2** |
| 27 | | $P_{50}$ | 1\|Genus/Species | 5 | 159.1 | 0 | |
| 28 | | $P_{50}$ + Forest | 1\|Genus/Species | 6 | 160.4 | 1.3 | |
| 29 |  | Forest | 1\|Genus/Species | 5 | 161.5 | 2.4 |  |
| 30 | | $P_{50}$ x Forest | 1\|Genus/Species | 7 | 163. | 3.8 | |

**Fig. 1.**
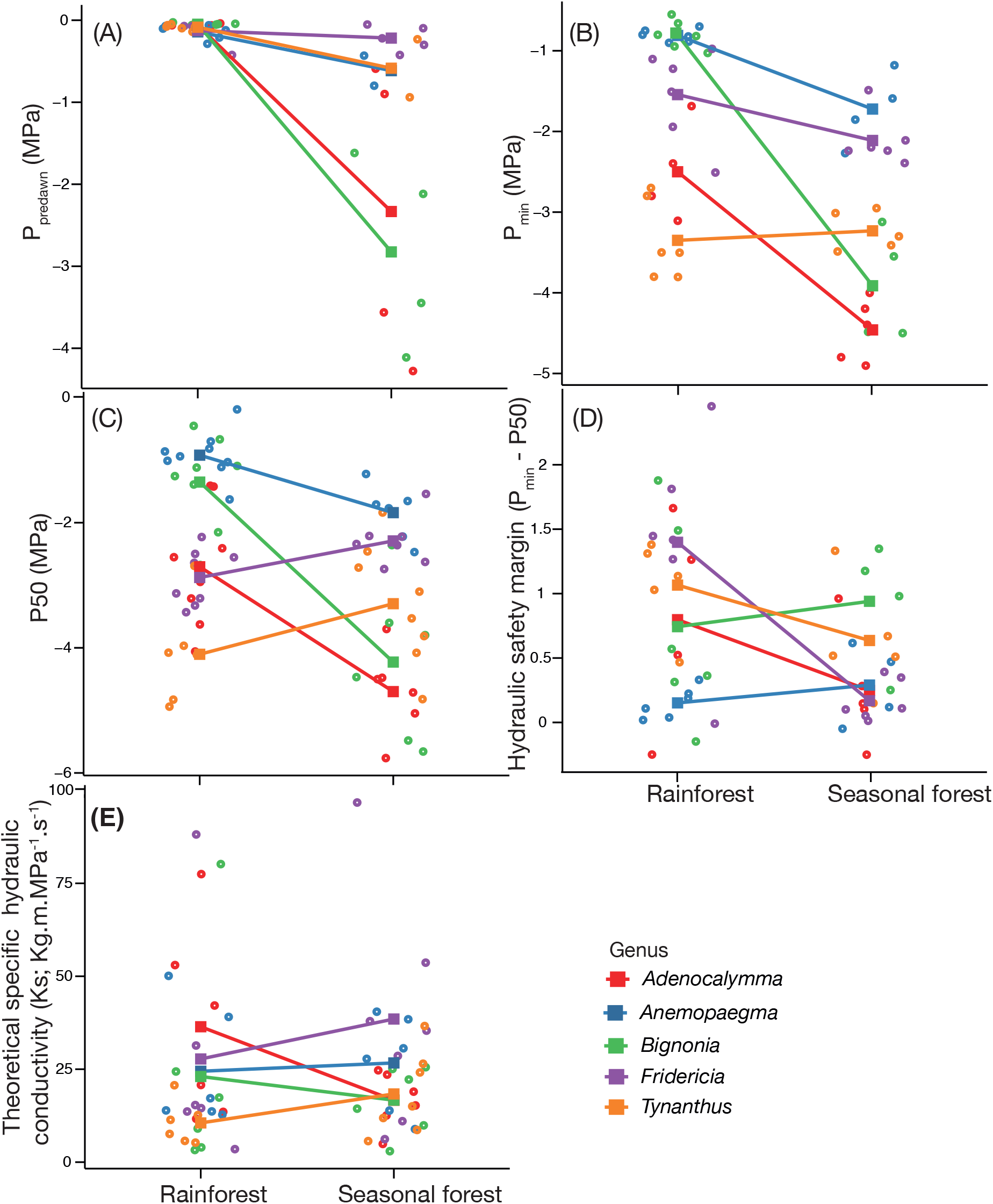
Comparison of hydraulic traits between two forest types across Bignonieae liana species: (A) Predawn water potential – P_predawn_, (B) Minimum seasonal water potential - P_min_, (C) hydraulic safety measured as the xylem pressure at which 50% of maximum amount of gas was discharged - P50, (D) hydraulic safety margin - HSM, and (E) theoretical specific hydraulic conductivity – Ks. We compared five pairs of congeneric liana species between two contrasting forest types: Amazon rainforest at Ducke Reserve and Atlantic seasonal forest at Santa Genebra Reserve. Lines connect values average (square) of pairs of congeneric species; colors indicate the five genera analyzed (red: *Adenocalymma;* blue: *Anemopaegma* green: *Bignonia;* purple: *Fridericia;* orange: *Tynanthus)*, and the open circles represent the hydraulic trait values per individual.

The hydraulic safety (P50) of liana species of the seasonal forest was 1.7 to 3.2 times more negative than liana species of the rainforest, although this difference was genus-dependent (Fig. 1C; Table 1; Model 11 in Table 2). Three congeneric pairs of Bignonieae lianas from the dry seasonal forest had higher hydraulic safety values (more negative P50) than the paired liana species of the rainforest. In contrast, the other two congeneric pairs, *Fridericia* and *Tynanthus*, had similar hydraulic safety values between the two forest types (Fig. 1C). The genus-dependent effect of forest type on hydraulic safety indicated that the higher hydraulic safety found in liana species of dry seasonal forest compared to liana species of wet rainforest repeatedly occurred at least in three phylogenetic lineages within the Bignonieae. The hydraulic safety margin (P_min_ – P50) was similar between liana species from the rainforest and seasonal forest (Model 16 in Table 2; Fig. 1D). The hydraulic safety margin ranged from −0.2 to 2 MPa in liana species of the rainforest, and from −0.2 to 1.5 MPa in liana species of the seasonal forest (Table 1; Fig. 1D).

The hydraulic efficiency as quantified by the theoretical specific conductivity (Ks) was similar between liana species from the rainforest and seasonal forest (Model 21 in Table 2; Fig. 1E). Ks was not related to hydraulic safety (P50) (Model 26 in Table 2, Fig. 2).

**Fig. 2.**
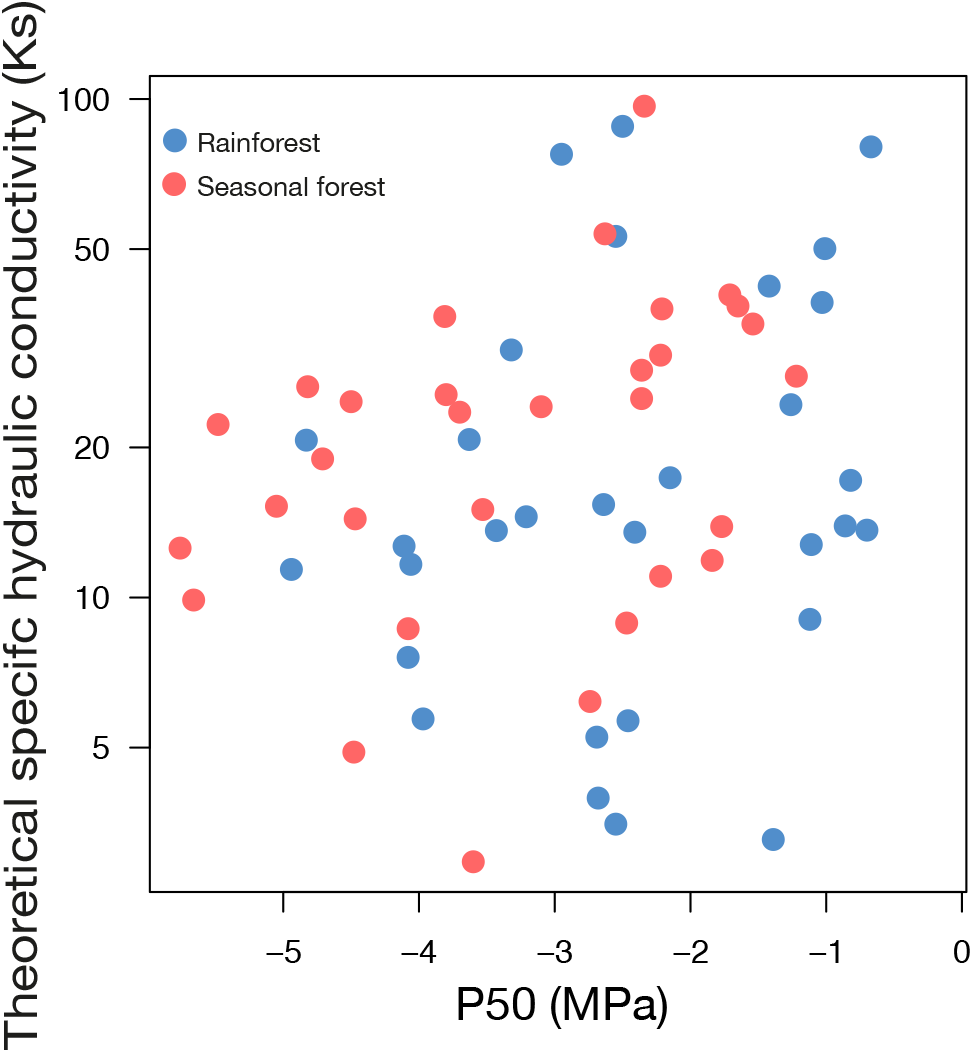
Relationship between theoretical specific hydraulic conductivity (Ks; on a log scale) and hydraulic safety (P50) across Bignonieae liana individuals in the rainforest (blue) and seasonal forest (red). In this analysis, we included 30 liana individuals of each forest type.

### Hydraulic safety of trees, shrubs, and lianas in tropical forests

Using the combined dataset (i.e. our data and literature), there was no difference in xylem resistance to embolism between liana species of rainforest and seasonal forests (t-test = −0.05; P = 0.95; Fig. 3). The average P50 of Bignonieae lianas from the rainforest was 10% more negative than the global average for tree and shrub angiosperms from the tropical rainforest (−2.01 MPa) (Fig. 3), but this difference was not significant (t-test = −0.81; P = 0.41). The average P50 of Bignonieae lianas from the seasonal forest was 21% more negative than the global average for tree and shrub angiosperms from the seasonal tropical forest (−2.58 MPa) (Fig. 3), but this difference also was not significant (t-test = 1.69; P = 0.09). The local variation in P50 for Bignonieae lianas (−3.18 and −2.86 MPa ranges in the rainforest and seasonal forest, respectively) covered about 30 % of the global variation in P50 for angiosperms (−10.9 MPa, ranging from −0.1 to −11 MPa, dataset indicated in the M&M section, Fig. 3). The variation in P50 for all lianas (−3.82 and −4.5 MPa for rainforest and seasonal forest, respectively) covered about 42% of the global variation in P50 for angiosperms (Fig. 3).

**Fig. 3.**
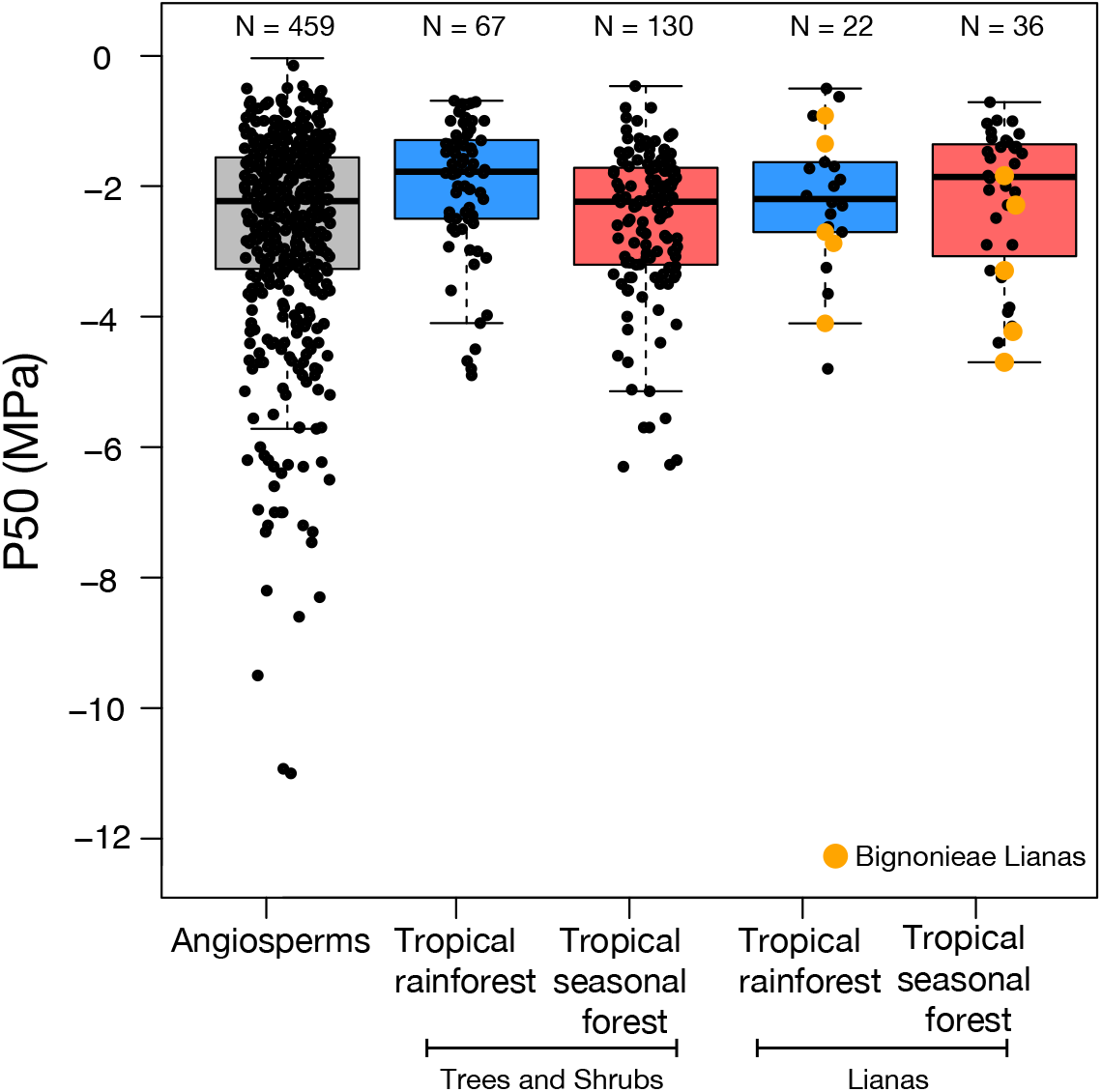
Boxplot of P50 (MPa; i.e., xylem pressure inducing 50% loss of hydraulic conductivity or at which 50% of maximum gas amount discharged) for angiosperm species, trees and shrubs, and lianas (Bignonieae lianas in orange) across tropical rainforest (blue) and seasonal tropical forest (red). Open circles were the average of P50 values per species for all angiosperm trees, shrubs, succulents, herbs, and lianas available in the global meta-data (dataset from: Zhu and Cao, 2009; Choat *et al.*, 2012; Carvalho *et al.*, 2016; Chen *et al.*, 2017, 2021; Oliveira *et al.*, 2019; Tan *et al.*, 2020; De Guzman *et al.*, 2021; Medina-Vega *et al.*, 2021; Smith-Martin *et al.*, 2022). Orange dots were average values of P50 of the five congeneric lianas species from the Amazon rainforest at Ducke Reserve (Amazonas, Brazil) and the seasonal forest at Santa Genebra Reserve (São Paulo, Brazil).

## Discussion

We investigated the hydraulic performance of lianas in Neotropical forests, more specifically, whether pairs of congeneric species from the tribe Bignonieae, one of the most representative and diverse clades of lianas in the Neotropics, have differences in safety and efficiency of water transport associated with climate seasonality. The first hypothesis was partially supported: lianas of the seasonal forest cope with a lower availability of water, have a higher hydraulic safety thus minimizing the risk of hydraulic failure (positive HSM) under drought, although this was not universal across all genera. Our second hypothesis was also partially supported: lianas species of both forest types invest in an equally efficient xylem in water transport. Therefore, although the liana species from the seasonal forest were not more efficient than lianas from the rainforest, we did not detect the hydraulic safety-efficiency trade-off. Furthermore, the increased hydraulic safety of liana species from the seasonal forest occurred at least three times within Bignonieae and indicates an adaptive drought convergence. Lastly, our third hypothesis was not supported: lianas of seasonal forests worldwide do not show higher embolism resistance, and in general, hydraulic safety is similar among lianas, shrubs, and trees from wet and seasonally dry tropical forests.

In sum, our results highlight a probable convergence in the hydraulic responses of the tribe Bignonieae living in seasonal forests and the equally high water-conducting efficiency across liana species as a general pattern of the lianescent life form. The similar positive safety margin and hydraulic efficiency suggest an analogous potential for survival and growth of lianas across tropical forests. Theses hydraulic features associated with an opportunistic strategy of rapid growth, especially in disturbed areas (Putz, 1984; Schnitzer *et al.*, 2005, 2015; Schnitzer, 2018) can favor the abundant liana species in seasonal forests, as observed for some Bignonieae liana species.

### Higher embolism resistance and similar conductive efficiency in lianas across tropical forest types

Plants from drier environments cope with lower water availability, reaching more negative seasonal minimum water potentials (Choat *et al.*, 2012, 2018; Mantova *et al.*, 2021; Oliveira *et al.*, 2021) as also observed on our seasonal forest species. The establishment and survival of plants in seasonally dry environments are related to drought resistance mechanisms (Choat *et al.*, 2008, 2010; Trifilò *et al.*, 2014; Santiago *et al.*, 2016). Greater drought resistance allows plants to operate at more negative xylem water potential with reduced levels of embolism within xylem (Gleason *et al.*, 2016). We found a higher embolism resistance (more negative P50) in three lineages of Bignonieae lianas from seasonal forests in comparison to rainforests, in agreement with findings for trees, supporting the perspective that drought tolerance linked to higher xylem resistance to embolism is a mechanism to cope with water scarcity and allow survival in drier environments across life-forms (Anderegg *et al.*, 2016; Barros *et al.*, 2019).

However, two lineages of Bignonieae did not have a higher hydraulic safety in the seasonal as compared to rainforests. The *Tynanthus* lineage had above average hydraulic safety values, even if this did not differ among forest types. The low hydraulic safety in the *Fridericia* lineage suggest that these species may invest in drought avoidance mechanisms to ensure the survival (Carvalho *et al.*, 2016). There is evidence that some liana species from arid environments and seasonal forests (e.g. *Fridericia)* have rapid stomatal control and leaf shedding, which can reduce the water loss as well as the risk of embolism under drought conditions (Carvalho *et al.*, 2016; Chen *et al.*, 2017). In addition, some species can have deep roots and capture water from deeper layers in the soil (Holbrook and Putz, 1996; Restom and Nepstad, 2001; Brum *et al.*, 2019), although the evidence for deep roots in lianas is still scarce (De Deurwaerder *et al.*, 2018; Smith-Martin *et al.*, 2019). Therefore, some tropical forest species with lower hydraulic safety may invest in more in avoidance than tolerance strategies to ensure survival (Oliveira *et al.*, 2021).

In the Amazon rainforest, given the high total annual rainfall and the short dry season, water availability is higher, and consequently, the less negative values of P_min_ and P50 found here were expected. However, xylem resistance to embolism was highly variable across liana species at the small spatial scale of a single Amazon forest. This high variability of P50 may be due to local differences in water availability along the hydro-edaphic gradient, which was associated to the differences in hydraulic resistance across tree species (Oliveira *et al.*, 2019). Similarly, we found the lianas with greater resistance to embolism farther away from water courses, i.e., in relatively drier conditions compared to those located in stream valleys. The coexistence of these lianas spatially close in the same forest is evidence of habitat partitioning associated with hydraulic features (Cosme *et al.*, 2017; Oliveira *et al.*, 2019). The higher hydraulic vulnerability of tree and liana species associated to valleys may be counterbalanced by the valley’ s wetter conditions, where groundwater supplies the soil all the time during ‘normal’ climatic years. However, these species may be at particular risk if the water-table level deepens during extreme droughts (Costa *et al.*, 2023).

The similar and positive hydraulic safety margin of our ten species suggests that their xylem does not experience a high risk of embolism formation, similar to findings for central and eastern rainforest trees (Brum *et al.*, 2019; Garcia *et al.*, 2021). However, the difference in HSM between lianas species within each forest type indicates that some species do have a greater risk of hydraulic failure. Damage to the xylem system with increased water stress reduces the conductive capacity, but can be relieved by producing new vascular tissue and/or tissue rehydration (Brodersen and McElrone, 2013; Knipfer *et al.*, 2016). There is evidence of cambial activity producing new vascular tissues during the wet season and early dry season in lianas of Bignonieae (Lima *et al.*, 2010; Angyalossy *et al.*, 2015) and Fabaceae (Brandes *et al.*, 2011), and throughout the year in Sapindaceae (León-Gómez and Monroy-Ata, 2005; Brandes *et al.*, 2022). Even if the damage to the hydraulic system of lianas is relatively high for some species, the small area of non-embolized xylem and water reserves in the parenchymal tissue must be sufficient to supply the young branches and leaves with water, avoiding desiccation and death of apical branches (Morris, 2016; Mantova *et al.*,2021).

The high and efficient water transport system in lianas has been shown in many studies (Carlquist, 1985; Ewers and Fisher, 1989; Ewers *et al.*, 1990). Lianas may develop a water transport system ten times more efficient than trees and shrubs (Gartner *et al.*, 1990; Gerolamo and Angyalossy, 2017; Zhu *et al.*, 2017). However, we found that Bignonieae lianas from both forest types do not differ in theoretical specific conductivity. Indeed, Medina-Vega *et al*. (2021) found no differences in the average xylem-specific conductivity between 16 other species from dry and wet forests. Unlike our initial hypothesis that seasonal forest lianas would be more efficient in transporting water, our result of similar efficient use of space for transporting water between forest types supports a more general pattern for lianas’ stems - an efficient hydraulic transport system with low mechanical demand for support.

The high-water transport efficiency of Bignonieae lianas is not related to hydraulic safety, as predicted by the safety-efficiency trade-off. Other lianas from different tropical forests also did not follow this trade-off (Zhu and Cao 2009; Van der Sande *et al.*, 2019; Medina-Vega *et al.*, 2021). Generally, the efficiency-safety trade-off has been weak or non-existing, regardless of plant habit (Gleason *et al.*, 2016; Van der Sande *et al.*, 2019). This decoupling of hydraulic efficiency from safety has recently been suggested based on a three-dimensional pit membrane model (Kaack *et al.*, 2021; Lens *et al.*, 2022).

### Embolism resistance is the result of convergent evolution

Drought survival strategies, such as hydraulic safety, may vary among populations, species, and lineages (Sperry and Saliedra, 1994; Lens *et al.*, 2013; Anderegg *et al.*, 2016; Dias *et al.*, 2019), as a result of repeated evolutionary adaptation under similar environments (Fontes *et al.*, 2020; Guillemot *et al.*, 2022) or phylogenetic conservatism (Losos, 2008;

Skelton *et al.*, 2021). We showed that increased hydraulic safety in Bignonieae lianas occurred in unrelated taxa under similar environments (dry seasonal forest), with substantial variation within genus. Nonetheless, this simple correlation does not explain why particular features are favored in specific habitats and additional evidence is needed to support a non-stochastic mechanism for convergent evolution (Losos, 2008; Stayton, 2015). Hydraulic safety has an adaptive significance as a mechanism of drought tolerance in trees and shrubs of evergreen angiosperms (Choat *et al.*, 2012; Anderegg *et al.*, 2016). Furthermore, the biogeographic history of Bignonieae highlights more recent migrations of lianas to dry forests and savannas repeatedly in different lineages (Lohmann *et al.*, 2013). This historical pattern and functional evidence for hydraulic safety indicate that hydric selective pressures could lead to the convergent evolutionary hydraulic specialization. However, the convergent pattern for hydraulic safety is not applicable for rainforest lianas because the biogeographic history indicates that these lianas have probably been under greater humidity throughout history. Therefore, the strong association between plant traits and each forest type implies that only the higher hydraulic safety of liana species detected in three lineages (genera) on the seasonal forest can be interpreted as an adaptation of lianas for restrictive hydric conditions.

### Embolism resistance of lianas in a global context

Based on our Bignonieae data, some liana species of seasonal forest have a high resistance to embolism as a strategy to deal with droughts. However, by combining hydraulic safety global data from other liana species collected in tropical forests of Panama and China, there is a high variability of resistance to xylem embolism and no difference between seasonally dry forests and rainforests. Also, the hydraulic safety is similar to those found in the other habits (Fig. 3). If these hydraulic safety values are not overestimated by technical artifacts as open-vessels for plants with long vessels (Wheeler *et al.*, 2013; Torres-Ruiz *et al.*, 2014; 2017; Pereira *et al.*, 2021), these results support that lianas have different strategies to deal with drought (Schnitzer *et al.* 2018) and that hydraulic safety is one of them.

Many lianas have a multifocal growing strategy with hydraulic redistribution and high water storage capacity (Morris *et al.*, 2016; de Azevedo Amorim *et al.*, 2018). If a branch dies due to mechanical damage or desiccation, new resprouts associated with other rooting points and storage reserve tissues can keep the plant alive (Rocha *et al.*, 2020). Therefore, high hydraulic and photosynthetic efficiency, production of new vascular tissue even in the dry season, and different drought resistance strategies help us to explain why some liana species grow rapidly, and tend to become relatively more abundant towards seasonal tropical forests. The higher density (and density increase) of some liana species can be a transitory phenomenon under more seasonal tropical forests if droughts continue to increase in intensity, since the risk of drought-induced mortality is higher for fast-growing plants. These hydraulic strategies have been documented for the most abundant species, but little is known about the rare species of lianas - most of their diversity. New research investigating the lethal threshold of dehydration associated with abundant and rare plant species and hydraulic resistance strategies may bring insights to this question.

### Conclusion

In this study, we show that seasonal forest lianas can be more hydraulically safe than rainforest lianas, although this result is not true for all liana species. Lianas from both forest types have also similar and high hydraulic efficiency. The higher hydraulic safety found for some lineages in Bignonieae associated with the evolutionary history of the group indicates an adaptive convergent evolution in seasonal forests. However, the global data of wood plants reviewed here indicated that lianas do not differ in hydraulic safety values compared to other habits in seasonal and wet tropical forests. The different hydraulic features found in this study associated with the multifocal growth strategy help us to understand, at least in part, the rapid growth of lianas in different tropical forests. Therefore, further studies are needed to explore, in addition to hydraulic safety, other functional strategies of lianas to deal with environmental water availability.

## Supplementary data

The following supplementary data are available at JXB online.

Fig. S1. Climate diagram of two contrasting tropical forests, with a wet (Ducke Reserve, Manaus, Brazil) and seasonally dry tropical forest (Santa Genebra Reserve, São Paulo, Campinas, Brazil).

Fig. S2. Hydraulic vulnerability curves for lianas Bignonieae from an Amazon rainforest at Ducke Reserve (Amazon) and a seasonal forest at Santa Genebra Reserve (Atlantic forest, São Paulo, Brazil).

Table S1. One voucher for each species was deposited at the University of São Paulo Herbarium (SPF; acronym according to Thiers, 2017).

Table S2. Large dataset used with average hydraulic safety (P50) for angiosperm species, including trees, shrubs, succulents, herbs and lianas from different forest sites.

## Acknowledgements

We thank Rafael Oliveira, Ellen Carvalho and Nara Vogado for her helpful suggestions. We acknowledge the invaluable help of Guilherme Antar, Valdiek Menezes, Elisangela Rocha, Lorena Bueno, Marilia Quinalha, Carlos Roberto and José Raimundo in the field and INPA’s (DSER) Department of Reserves for Logistical Support.

## Author contributions

CSG, AN, and VA designed research. CSG and AN collected the hydraulics data. CSG, AN, FRCC, LP and SJ analyzed the hydraulics data. CSG and AN wrote the first version of the manuscript with substantial input from FRCC, LP SJ and VA, and all authors contributed to revisions.

## Data availability

The average species trait data are available upon reasonable request from the authors.

